# A genomic footprint of a moving hybrid zone in marbled newts

**DOI:** 10.1101/2020.05.27.118448

**Authors:** Jan W. Arntzen, Julia López-Delgado, Isolde van Riemsdijk, Ben Wielstra

**Affiliations:** Naturalis Biodiversity Center, Leiden, The Netherlands; Institute of Biology, Leiden University, Leiden, The Netherlands; University of Leeds, Leeds, United Kingdom

**Keywords:** Amphibia, enclave, hybridization, Iberian Peninsula, Ion Torrent, introgression, species replacement, *Triturus marmoratus*, *Triturus pygmaeus*

## Abstract

We developed a panel of 44 nuclear genetic markers and applied this to two species of marbled newts in the north (*Triturus marmoratus*) and the south (*T. pygmaeus*) of the Iberian Peninsula, to document pattern and process of interspecific gene flow. The northernmost occurrence of *T. pygmaeus* genetic material was in a *T. marmoratus* population north of the Vouga river estuary. This suggested the past presence of a hybrid zone, possibly coinciding with a natural river outlet at ca. 1200 A.D. Since 1808, the species contact has moved back south to a by then completed, man-made Vouga channel. We also found evidence for a *T. marmoratus* genomic footprint in *T. pygmaeus* from the Serra de Sintra, near Lisbon. In combination with a previously reported southern, relic occurrence of *T. marmoratus* in between both areas, the data point to the superseding with hybridization of *T. marmoratus* by *T. pygmaeus*. We estimate that the species hybrid zone has moved over a distance of ca. 215 km.

## Introduction

Hybrid zones occur when the geographic ranges of closely related and reproductively incompletely isolated taxa meet and cross-fertilize to produce admixed offspring (Barton and Hewitt, 1985; Harrison, 1990). Hybrids typically show a considerable reduction in fitness relative to their parents. When dispersal into the contact zone fuels the hybridization and selection process, an equilibrium may arise, as in so-called ‘tension hybrid zones’ (Barton, 1979). This type of hybrid zone, where the antagonistic effects of dispersal of parental forms and selection against hybrids balance each other, typically shows a sigmoid (or ‘clinal’) genetic profile (Bazykin, 1969; Szymura and Barton, 1986). At points of initial contact, one of the two species may have a competitive advantage over the other and species replacement would entail hybrid zone movement (Barton, 1979; Buggs, 2007; Wielstra, 2019). Yet, because hybrid zones often settle at an ecotone (Endler, 1977; Kruuk et al., 1999; Bierne et al., 2011), or are captured at areas of low dispersal (Barton, 1979; Barton and Hewitt, 1985; Abbott et al., 2013), mosaic distributions can also be formed, with a spatial grain that depends on the ecological preference of the species in conjunction with the distribution of suitable habitats over the landscape.

Hybrid zone movement may be inferred from historical documentation or from the reconstruction of relic distribution patterns. Both approaches have, however, limited applicability. First, while historical records may be informative to recently formed and fast-moving hybrid zones, most hybrid zones are old and have formed hundreds or thousands of years ago (Hewitt, 1988, 2000). Second, pockets of persistence of one species enveloped by the other may be indicative of past species replacement. However, for such an enclave pattern to be formed crucially depends on the local distribution of differentially preferred habitat types, on a limited species dispersal capability and, possibly, on limited hybridization. Ground-dwelling organisms with low dispersal, such as many amphibian species, may be prime candidates to demonstrate enclave formation.

Hybrid zone movement may also be inferred from genetic data. In the wake of a moving hybrid zone, the introgression of neutral alleles is exaggerated from the retreating to the advancing species and, because this introgression is only subject to drift, it will be geographically stable on an evolutionary time scale (Barton and Hewitt 1985; Currat et al. 2008). This pattern of genome-wide asymmetric introgression has been coined a genetic (or genomic) footprint of hybrid zone movement (Scribner and Avise, 1993). The nine species of *Triturus* newts have shown to provide a useful system for the study of hybrid zone movement, for which Arntzen et al. (2014) and Wielstra et al. (2017ab) provide examples from across the genus’ range. We here report on the development and evaluation of a large panel of nuclear genetic markers for hybrid zone research in the three western European species *Triturus cristatus* (Laurenti, 1768) (northern crested newt), *T. marmoratus* (Latreille, 1800) (northern marbled newt) and *T. pygmaeus* Wolterstorff, 1905 (pygmy marbled newt).

In the west of France, isolated pockets of *T. marmoratus* persist in an area documented to have witnessed the widespread advance of, and replacement by *T. cristatus* (Arntzen and Wallis, 1991; Visser et al, 2017). Morphologically determined F1 hybrids were shown to occur at a frequency of ca. 4% and introgression was reported from ten diagnostic allozyme markers, at a frequency of 0.3 % (Vallée, 1959; Schoorl and Zuiderwijk, 1981; Arntzen and Wallis, 1991; see also Arntzen et al., 2009). Clearly, to further evaluate pattern and process of species replacement and hybrid zone movement in this system, access to an enlarged molecular toolkit would be an asset.

In the west of Portugal, we also observed *T. marmoratus* in a local pocket away from its main range (Espregueira Themudo and Arntzen, 2007a). Because *T. marmoratus* is locally entirely surrounded by *T. pygmaeus* and separated from its main distribution by a distance in excess of that covered by regular dispersal, this pocket qualifies as an enclave. Employing four partially diagnostic allozyme markers, Espregueira Themudo and Arntzen (2007a) estimated the frequency of introgressed individuals at 1.0%, in addition to one locality (Juncal, coordinates 39.60N, 8.92W) where both species were found along with a possible backcross hybrid. The presence of an area where *T. marmoratus* endured, but was intersected and excised from the main distribution by *T. pygmaeus*, led the authors to conclude that the latter species has been enlarging its range at the expense of the former species. Hybridization and introgression were limited, so that the genetic integrity of the species remained largely intact. Elsewhere along the species contact the species hybridization and introgression appear more frequent, with a clinal rather than a mosaic or bimodal species transition (Figure 1).

**Figure 1.**
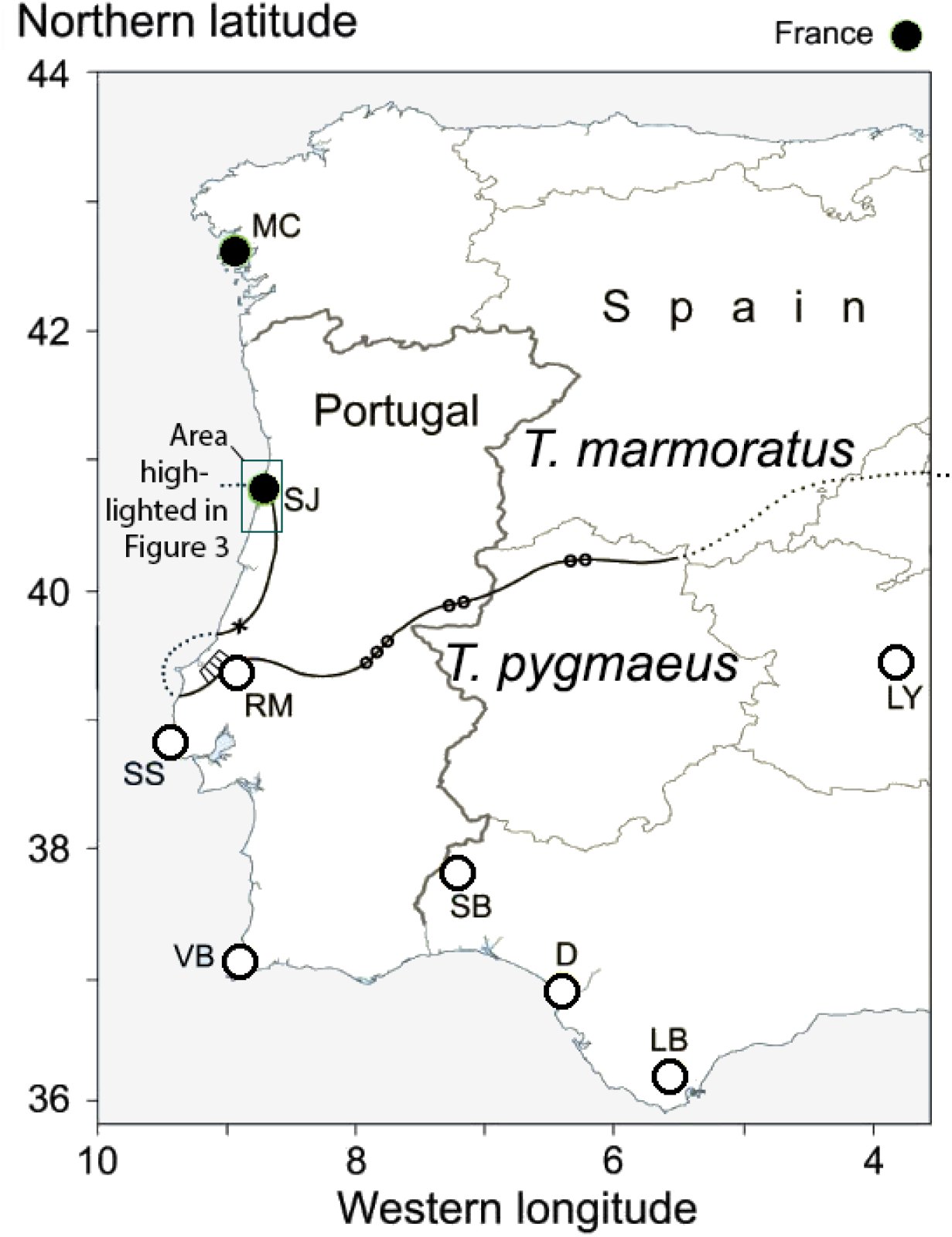
Approximate position of the abutting range borders of *Triturus marmoratus* and *T. pygmaeus* across the western part of the Iberian Peninsula. Solid and open large round symbols represent sampled populations of either species (Table 1). Populations tested for interspecific gene flow are SJ, RM and SS (details see text). Three areas where introgressive hybridization is well documented are marked by dots (after Arntzen, 2018) and one locality with both species is marked by a cross (population Juncal; Espregueira Themudo and Arntzen, 2007a). A *T. marmoratus* enclave, as more precisely documented in the latter study, is indicated by hatching (adjacent to locality RM).

We here employ 52 newly developed markers in the *T. marmoratus – T. pygmaeus* system in the west of Portugal with the aims i) to test that the species do hybridize, and ii) to search for the presence of a genetic remnants that were transferred to the superseding species. The key populations in our study are two potentially hybridized ones, just north and southeast of the mutual species border and one from the Serra de Sintra near Lisbon. We hypothesize the past presence of *T. marmoratus* in these mountains on account of i) its geographical position not far south (ca. 70 km) of the area of species replacement, and ii) the species’ documented ecological differentiation, with *T. marmoratus* prevailing at higher altitudes than *T. pygmaeus* (Arntzen and Espregueira Themudo, 2008). If *T. marmoratus* was present in the Lisbon peninsula, the population persisting for longest is likely to have been at the Serra de Sintra.

**Table 1.**
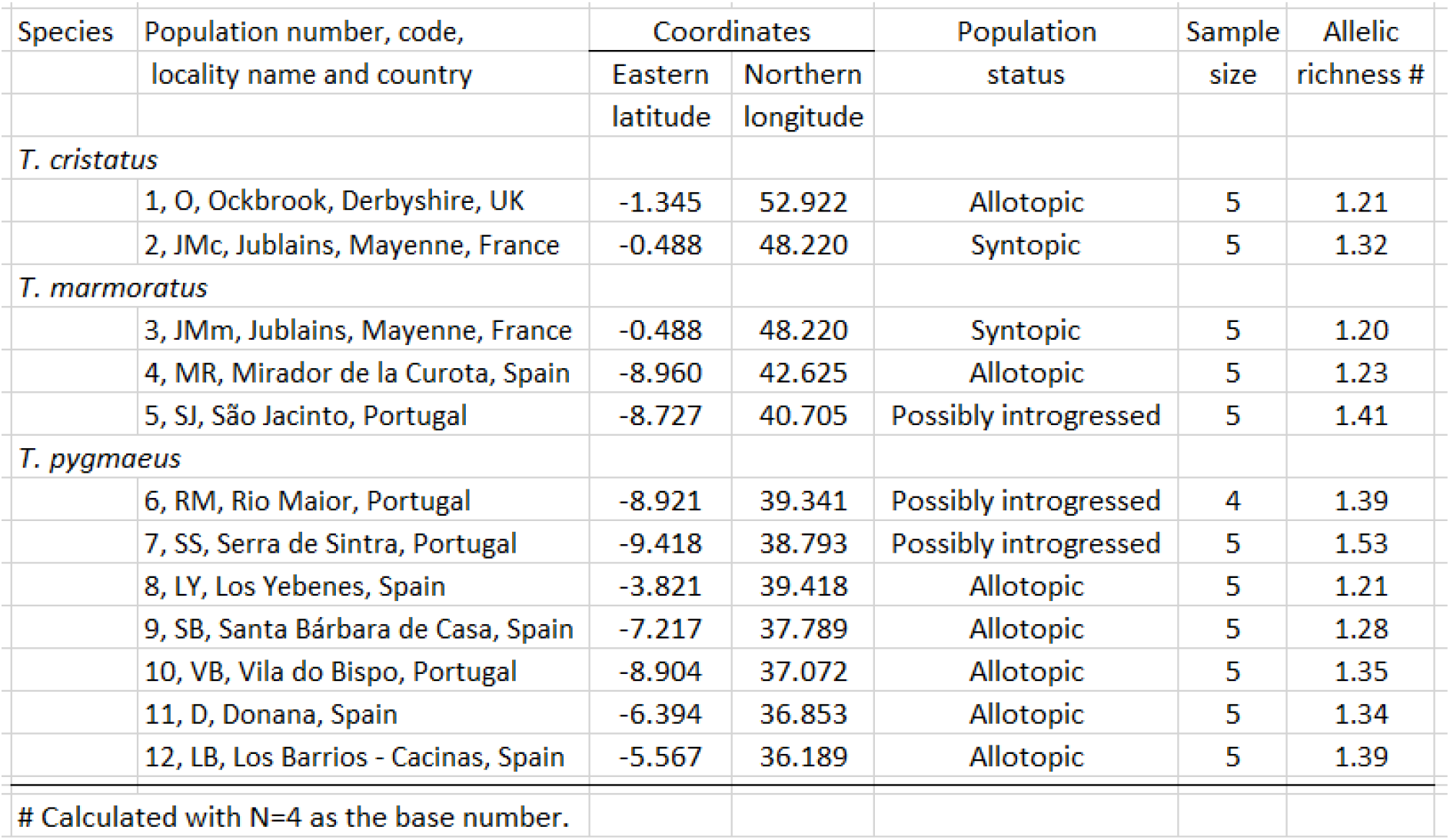
Populations of three species of *Triturus* newts subjected to genetic investigation.

## Materials and methods

We sampled five individuals per locality for seven allotopic and three potentially introgresssed populations of *T. marmoratus* and *T. pygmaeus.* We also studied *T. cristatus* in one population in allopatry and one in syntopy with *T. marmoratus*, all in western Europe (N=60, Figure 1, Table 1).

Using the Ion Torrent next-generation sequencing protocol described in Wielstra et al. (2014), we sequenced 52 nuclear markers. In brief, we amplified markers of ca. 140 bp in length (excluding primers), positioned in 3’ untranslated regions, in five multiplex PCRs. We pooled the multiplexes for each individual and ligated unique tags to be able to recognize the product belonging to each individual. We sequenced the amplicons on the Ion Torrent next-generation sequencing platform and processed the output with a bioinformatics pipeline that filters out poor quality reads and converts data to the FASTA format for further analysis. The total number of Ion Torrent reads after data filtering was 3,254,028. Mean coverage was 1,043 reads (range 0-18,749) per marker-individual combination and 93.8% of marker-individual combinations were considered successful (at least 20 reads per marker and per individual).

The sequences were realigned with AliView (Larsson, 2014) and subsequently converted into a genotypic data format by recoding the different allelic variants for each marker to a unique integer, resulting in two integers per individual per marker. Internal gaps were treated as informative insertion/deletion polymorphisms (indels). Data were taken out for eight loci that showed no variation or had more than five missing observations. Remaining missing data points amounted to N=56 across 44 loci and 60 individuals (2.1%). Haplotype networks were constructed with PopArt (Leigh and Bryant, 2015), which analysis confirmed the coherence of the data set, but also pointed to individual P_2826 from the Rio Major *T. pygmaeus* population as an outlier. The aberrant data point was removed from the data set. Basic population genetic analyses were carried out with GenePop (Rousset, 2008) and allelic richness (AR) was calculated with HP-Rare (Kalinowski, 2005). Population genetic differentiation was further analysed by principal component analysis (PCA) with Adegenet (Jombart, 2008).

## Results

The number of alleles observed per locus ranged from two to 16 and total numbers increased in the order *T. cristatus* – *T. marmoratus* – *T. pygmaeus*. Within the latter two species, allelic richness was lower in the seven allotopic populations (range 1.20-1.39) than in the three potentially introgressed populations São Jacinto, Rio Major and Serra de Sintra (range 1.39-1.53, Table 1). Population genetic differentiation ranged from *F*_st_=0.466 for *T. marmoratus* versus *T. pygmaeus* to *F*_st_=0.846 for *T. marmoratus* versus *T. cristatus*. The numbers of loci that contributed significantly to species discrimination (P<0.05 in Chi^2^ tests) were 20 out of 44 and 43 out of 44, respectively.

The first axis of a PCA widely separated *T. cristatus* from the other two species (results not shown). We found no indications of genetic admixture or introgression among syntopic *T. cristatus* and *T. marmoratus*. Further inspection shows the separation of *T. marmoratus* and *T. pygmaeus* along the second PCA-axis, with low or moderate levels of intraspecific variation (Figure 2). The *T. marmoratus* population of São Jacinto stands out as genetically differentiated from the other, more northern populations, with significant contributions by ten loci (P<0.05 in Chi^2^ tests). The overall level of differentiation is also significant (Fisher’s method, Chi^2^=111.5, df=44, P<0.001). For *T. pygmaeus*, no displacements towards *T. marmoratus* were observed, except for individual P_3576 from population Serra de Sintra (Figure 2). Along the third PCA-axis substantial genetic variation was observed within *T. pygmaeus* populations. Population Los Barrios was differentiated from the remainder (Fisher’s method, overall Chi^2^=234.7, df=64, P<0.001), with significant contributions by 13 loci (P<0.05 in Chi^2^ tests).

**Figure 2.**
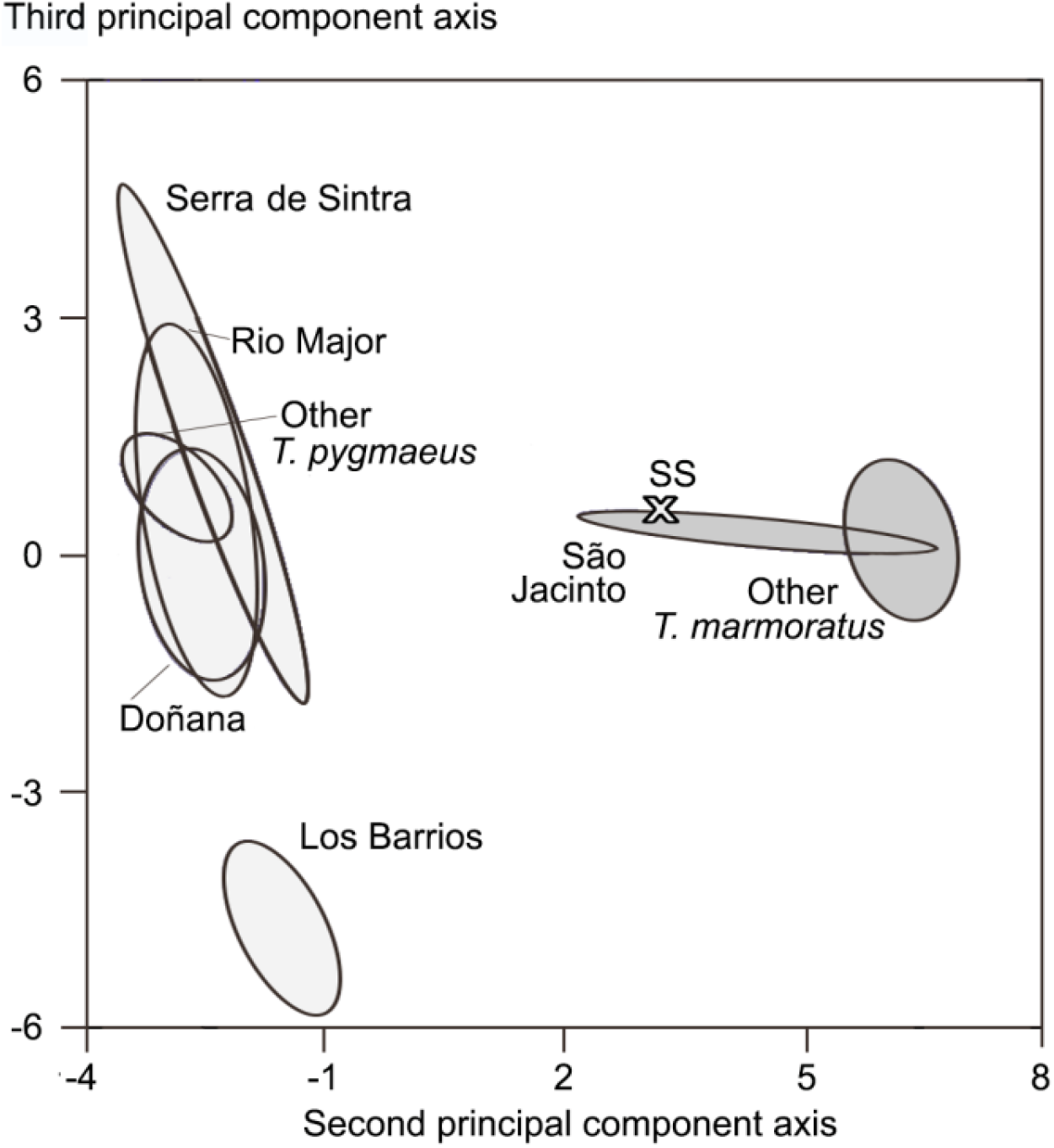
Results of a principal component analysis (PCA) of nuclear genetic data for *Triturus marmoratus* (dark shading) and *T. pygmaeus* (light shading). Data for populations are summarized by ellipses that represent the mean ± one standard deviation. Note that the inference of *T. marmoratus – T. pygmaeus* genetic admixture associates with the geographical position of these populations close to the mutual range border at either the present, as in São Jacinto, or possibly in the past, as hypothesized for Serra de Sintra (with one possibly admixed individual shown separately, SS). The third species in the study, *T. cristatus,* is deeply differentiated along the first PCA-axis (results not shown). The amount of the total variation explained along the first, second and third PCA-axis is 52.5%, 17.6% and 3.5%, respectively.

## Discussion

We developed a large panel of nuclear genetic markers and applied this to two species pairs of *Triturus* newts in the west of Europe, to scrutinize marker diagnosticity and to test for possible interspecific gene flow. *Triturus cristatus* and *T. marmoratus* are morphologically and genetically deeply differentiated. In spite of their differences, the species do hybridize in the area of range overlap in the west of France with, however, exceedingly low levels of introgression (Arntzen and Wallis, 1991). We were unable to confirm introgression, which is best attributed to the small sample sizes involved. We suggest that a selection of our high-performance diagnostic markers should be applied to a wider sample collected in the area of sympatry.

*Triturus marmoratus* and *T. pygmaeus* are similar, yet morphologically diagnosable species (García-París et al., 1993; Arntzen, 2018). They are genetically distinct (Espregueira Themudo and Arntzen, 2007b; Wielstra et al., 2019), with genetic admixture reported over several sections of their parapatric species border (Figure 1). We further investigated the Atlantic part of the range border with three targeted populations close to the present day (São Jacinto and Rio Major) and hypothesized past mutual range border (Serra de Sintra).

The distribution of *T. pygmaeus* currently reaches along the coast northwards up to the Aveiro Lagoon (coordinates 40.65N, 8.73W) that connects the river Vouga to the Atlantic Ocean. The presence of an enclave further south suggests that *T. pygmaeus* has been enlarging its range at the expense of *T. marmoratus*. The population of São Jacinto is genetically differentiated from other *T. marmoratus* populations and appears more similar to *T. pygmaeus*. These results suggest hybridization and introgression, but how to explain the presence of *T. pygmaeus* genes in the São Jacinto spit of land ? Colonisation from the east, the west or the north can be excluded because this involves crossing the lagoon, the ocean, or a long stretch of *T. marmoratus* territory. Recent dispersal from the south is precluded by the current Vouga discharge, the ca. 280 m wide ‘Barra Nova’. However, prior to this man-made outlet, the Vouga had its Atlantic discharge at a series of different, natural positions, ever since the estuary was formed in medieval times (Dias et al., 2012). If *T. pygmaeus* arrived early, its northerly advance may have been halted by one of the original Vouga channels north of São Jacinto (positions A and B in Figure 3). In case of a later arrival, five more southerly located river outlets (positions C, D, E, F and G) that were in place from ca. 1500 to 1757 may have impeded, but would not have stopped its northerly advance. If the *T. pygmaeus* – *T. marmoratus* hybrid zone had a southerly position, it must have been fairly wide for the Barra Nova to cut off a substantial part of its northern tail. This is a little unlikely because elsewhere in Portugal and Spain the hybrid zone for nuclear genetic markers is narrow (Arntzen, 2018), if not abrupt (Espregueira Themudo and Arntzen, 2007a). If the hybrid zone was positioned north of the Barra Nova, e.g. coinciding with former Vouga channels at positions A or B, entire *T. pygmaeus* populations may have been cut off from the main species range. This might have allowed the further northward advance of *T. pygmaeus*, but this scenario is not supported by the data. An alternative explanation is that the hybrid zone got attracted to a position were dispersal is low, such as across a waterway. This explanation we prefer, because it is in line with hybrid zone theory. Accordingly, we propose that the arrival of new recruits at São Jacinto, required to feed the northward expansion of *T. pygmaeus*, got truncated ever since the Barra Nova channel was completed in 1808 and that instead *T. marmoratus* regained the southern tip of the São Jacinto peninsula.

**Figure 3.**
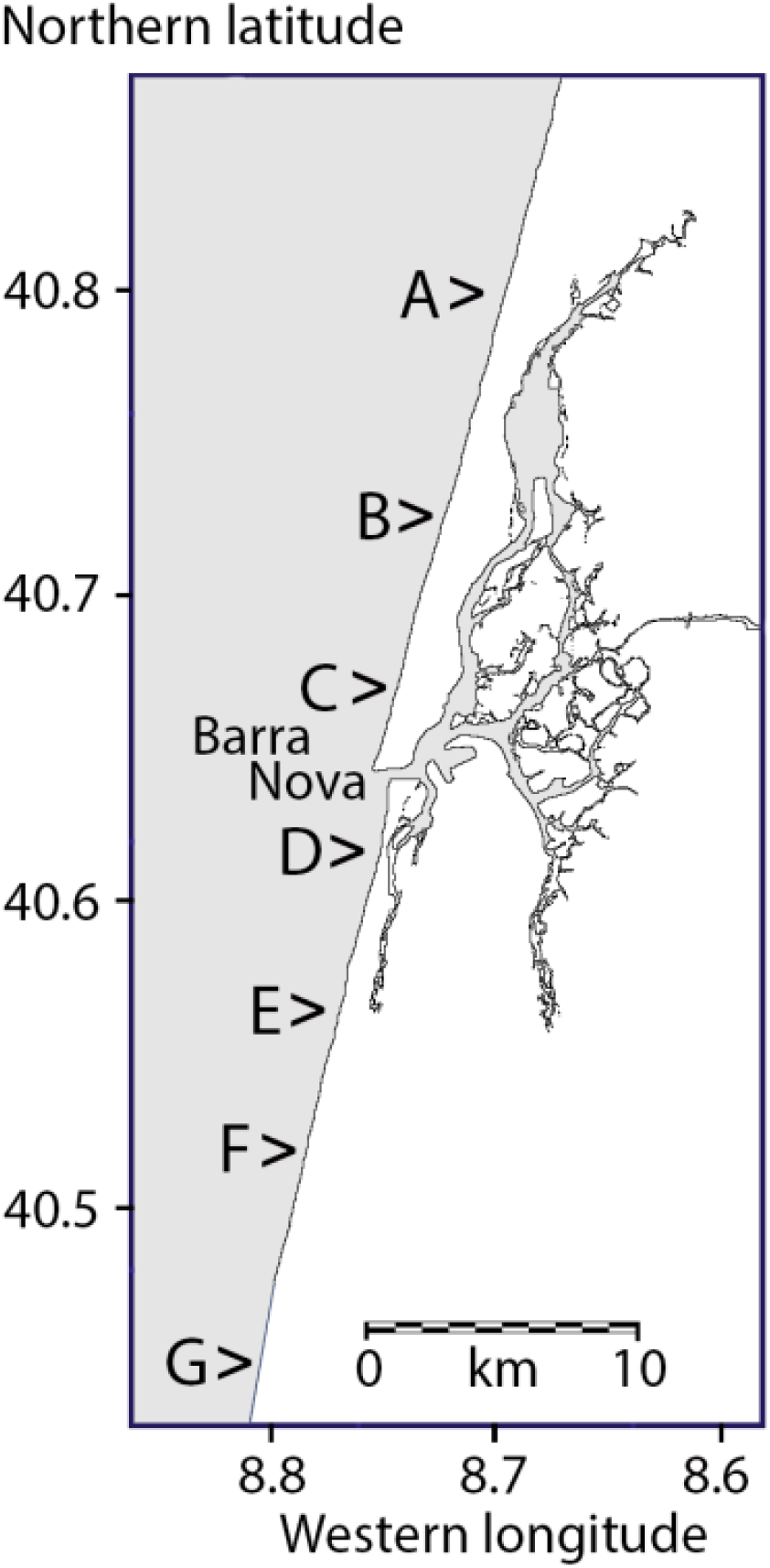
Western Portugal with the Aveiro Lagoon and the Atlantic coastline at the present day, with the discharge of the Rio Vouga at the Barra Nova. The Barra Nova is a man-made channel that was completed in 1808. Earlier, natural positions of the mouth of the Vouga were at positions A – 12^th^ century, B – ca. 1200, C – ca. 1500, D – 1584, E – 1643, F – 1685 and G – 1757. Redrawn from Dias et al. (2012). The study locality São Jacinto is located in between B and C. For the wider position of the area see Figure 1.

The *T. pygmaeus* population at Rio Major shows no signal of genetic admixture or introgression. The nearest genetically investigated *T. marmoratus* populations are at Salir de Matos and Fonte da Pena da Gouvinha (Arntzen, 2018) at distances of ca. 17 km. This exceeds the observed (Kupfer, 1998; Kupfer and Kneitz, 2000; Trochet et al., 2014) or inferred (Arntzen and Wallis, 1991; Wielstra et al., 2017b) dispersal capability of *Triturus* newts by an order of magnitude. To further our understanding, a more detailed survey is required to document position, shape and width of the hybrid zone in this area.

A remarkable feature of the *T. pygmaeus* population of the Serra de Sintra is its high level of genetic variability that is expressed along the third PCA-axis (Figure 2). Our sample also includes one individual that carries a complement of genetic material typical for *T. marmoratus*. Because *T. marmoratus* is unknown for the region, we interpret the signal as a genetic footprint derived from the species’ past presence in a distribution relic that by now has all but vanished. Similarly, the footprint can be seen as ‘islands of northern alleles’ (cf. Macholán et al., 2011). This explanation conforms to the hypothesis of species replacement with hybridization that instigated our study. Alternatively, alleles shared with *T. marmoratus* may reflect incomplete lineage sorting, but it is unclear why this would apply to the Serra de Sintra population and not to reference populations.

Finally, the southernmost studied *T. pygmaeus* population of Los Barrios is genetically distinct from northern ones. This observation parallels that on other amphibian genera with taxa endemic to the Betic region, including fire salamanders (in casu *Salamandra salamandra longirostris* Joger and Steinfartz, 1994) and midwife toads (in casu *Alytes dickhilleni* Arntzen and García-Paris, 1995). The range border of the southern potential lineage of *T. pygmaeus* has yet to be determined, but could involve the Guadalquivir river.

## Concluding remarks

In France *T. cristatus* and *T. marmoratus* have overlapping ranges whereas in Spain and Portugal *T. marmoratus* and *T. pygmaeus* keep a parapatric range border. Sympatry as observed in the former species pair may be associated with a low level of interspecific geneflow (Barton and Hewitt, 1985). Their overall genetic integrity is accounted for by post-mating isolating factors such as genetic incompatibility (Arntzen and Wallis, 1991; Arntzen et al., 2009), and pre-mating isolating factors including different habitat preferences (Schoorl and Zuiderwijk, 1981; Visser et al., 2017) or possibly differences in mating behaviour (Zuiderwijk, 1990). Conversely, the mutual ranges of *T. marmoratus* and *T. pygmaeus* are contiguous, with more or less frequent hybridization and introgression that is restricted to a narrow and clinal transition zone (Arntzen, 2018). However, along the Atlantic coast the zone shows aspects of a mosaic, in particular around Caldas da Rainha (coordinates 39.41N, 9.14W), where a pocket of six or more *T. marmoratus* populations is surrounded by *T. pygmaeus* (Espregueira Themudo and Arntzen, 2007a). The documented enclave forms the strongest evidence so far for local species replacement of *T. marmoratus* by *T. pygmaeus.* A *T. marmoratus* genetic footprint at Serra de Sintra, as suggested by our new data, is in line with this interpretation. We pinpointed the northern fringe of the hybrid zone at the Vouga and the distance over which it appears to have moved, from the Serra de Sintra, over Caldas to the Barra Nova, amounts to 215 km.

## Acknowledgments

We thank Eva Albert, Rudolf Malkmus, Iñigo Martínez-Solano and Rui Rebelo for locality information or help with tissue sampling.

## References

Abbott, R., Albach, D., Ansell, S., Arntzen, J. W., Baird, S. J., Bierne, N., Boughman, J., Brelsford, A., Buerkle, C. A., Buggs, R., Butlin, R. K., Dieckmann, U., Eroukhmanoff, F., Grill, A., Cahan, S. H., Hermansen, J. S. G., Hewitt, G, Hudson, A. G., Jiggins, C., Jones, J., Keller, B., Marczewski, T., Mallet, J., Martinez Rodriguez, P., Möst, M., Mullen, S., Nichols, R., Nolte, A. W., Parisod, C., Pfennig, K., Rice, A. M., Ritchie, M. G., Seifert, B., Smadja, C. M., Stelkens, R., Szymura, J. M., Väinölä, R., Wolf, J. B. W. & Zinner, D. (2013). Hybridization and speciation. Journal of Evolutionary Biology, 26, 229–246. https://doi.org/10.1111/j.1420-9101.2012.02599.x

Anderson, E. C., & Thompson, E. A. (2002). A model-based method for identifying species hybrids using multilocus genetic data. Genetics, 160, 1217–1229.

Arntzen, J. W. (2018). Morphological and molecular characters to describe a marbled newt hybrid zone in the Iberian peninsula. Contributions to Zoology, 87, 167–185. https://doi.org/10.1163/18759866-08703003

Arntzen, J. W., & Espregueira Themudo, G. (2008). Environmental parameters that determine species geographical range limits as a matter of time and space. Journal of Biogeography, 35, 1177–1186. https://doi.org/10.1111/j.1365-2699.2007.01875.x

Arntzen, J. W., & Wallis, G. P. (1991). Restricted gene flow in a moving hybrid zone of the newts *Triturus cristatus* and *T. marmoratus* in western France. Evolution, 45, 805–826. https://doi.org/10.1111/j.1558-5646.1991.tb04352.x

Arntzen, J. W., Jehle, R., Bardakci, F., Burke, T., & Wallis, G.P. (2009). Asymmetric viability of reciprocal-cross hybrids between crested and marbled newts (*Triturus cristatus* and *T. marmoratus*). Evolution, 63, 1191–1202. https://doi.org/10.1111/j.1558-5646.2009.00611.x

Arntzen, J. W., Wielstra, B., & Wallis, G. P. (2014). The modality of nine *Triturus* newt hybrid zones assessed with nuclear, mitochondrial and morphological data. Biological Journal of the Linnean Society, 113, 604–622. https://doi.org/10.1111/bij.12358

Barton, N. H. (1979). The dynamics of hybrid zones. Heredity, 43, 341–359. https://doi.org/10.1038/hdy.1979.87

Barton, N. H., & Hewitt, G. M. (1985). Analysis of hybrid zones. Annual Review of Ecology and Systematics, 16, 113–148. https://doi.org/10.1146/annurev.es.16.110185.000553

Bazykin, A. D. (1969). Hypothetical mechanism of speciation. Evolution, 23, 685–687. https://doi.org/10.1111/j.1558-5646.1969.tb03550.x

Bierne, N., Welch, J., Loire, E., Bonhomme, F., & David, P. (2011). The coupling hypothesis: Why genome scans may fail to map local adaptation genes. Molecular Ecology, 20, 2044–2072. https://doi.org/10.1111/j.1365-294X.2011.05080.x

Buggs, R. J. A. (2007). Empirical study of hybrid zone movement. Heredity, 99, 301–312. https://doi.org/10.1038/sj.hdy.6800997

Currat, M., Ruedi, M., Petit, R. J., & Excoffier, L. (2008). The hidden side of invasions: Massive introgression by local genes. Evolution, 62, 1908–1920. https://doi.org/10.1111/j.1558-5646.2008.00413.x

Dias, J. A., Bastos, M. R., Bernardes, C., Freitas, J. G., & Martins, V. (2012). Interacções Homem-Meio em zonas costeiras: o caso de Aveiro, Portugal. In Baía de Sepetiba – Estado da Arte. Eds M. Antonieta, C. Rodrigues, S. Dias Pereira & S. J. Barbosa dos Santos (Pp. 215–235). Rio de Janeiro, Brazil: Editora Corbã.

Endler, J. A. (1977). Geographic variation, speciation, and clines. Princeton, NJ: Princeton University Press.

Espregueira Themudo, G., & Arntzen, J. W. (2007a). Newts under siege: range expansion of *Triturus* pygmaeus isolates populations of its sister species. Diversity and Distributions, 13, 580–586 https://doi.org/10.1111/j.1472-4642.2007.00373.x

Espregueira Themudo, G., & Arntzen, J. W. (2007b). Molecular identification of marbled newts and a justification of species status for *Triturus marmoratus* and *T. pygmaeus*. Herpetological Journal, 17, 24–30.

García-París, M., Herrero, P., Martín, C., Dorda, J., Esteban, M., & Arano, B. (1993). Morphological characterization, cytogenetic analysis, and geographical distribution of the Pygmy marbled newt *Triturus marmoratus pygmaeus* (Wolterstorff, 1905) (Caudata: Salamandridae). Contributions to Zoology, 63, 3–14. https://doi.org/10.1163/26660644-06301001

Harrison, R. G. (1990). Hybrid zones: windows on evolutionary process. Oxford Surveys in Evolutionary Biology, 7, 69–128.

Hewitt, G. M. (1988). Hybrid zones – natural laboratories for evolutionary studies. Trends in Ecology and Evolution, 3, 158–167. https://doi.org/10.1016/0169-5347(88)90033-X

Hewitt, G. M. (2000). The genetic legacy of the Quaternary ice ages. Nature, 405, 907–913. https://doi.org/10.1038/35016000

Jombart, T. (2008). Adegenet: a R package for the multivariate analysis of genetic markers. Bioinformatics, 24, 1403–1405. https://doi.org/10.1093/bioinformatics/btn129

Kalinowski, S. T. (2005). HP-Rare: a computer program for performing rarefaction on measures of allelic diversity. Molecular Ecology Notes, 5, 187–189. https://doi.org/10.1111/j.1471-8286.2004.00845.x

Kruuk, L. E. B., Baird, S. J. E., Gale, K. S., & Barton, N. H. (1999). A comparison of multilocus clines maintained by environmental adaptation or by selection against hybrids. Genetics, 153, 1959–1971.

Kupfer, A. (1998). Wanderstrecken einzelner Kammolche (*Triturus cristatus*) in einem Agrarlebensraum. Zeitschrift für Feldherpetologie, 5, 238–241.

Kupfer, A., & Kneitz, S. (2000). Population ecology of the great crested newt (*Triturus cristatus*) in an agricultural landscape: dynamics, pond fidelity and dispersal. Herpetological Journal, 10, 165–171.

Larsson, A. (2014). AliView: a fast and lightweight alignment viewer and editor for large data sets. Bioinformatics, 30, 3276–3278. https://doi.org/10.1093/bioinformatics/btu531

Leigh, J. W., & Bryant, D. (2015). PopART: Full-feature software for haplotype network construction. Methods in Ecology and Evolution, 6, 1110–1116. https://doi.org/10.1111/2041-210X.12410

Macholán, M., Baird, S.J., Dufková, P., Munclinger, P., Bímová, B.V., & Piálek, J. (2011). Assessing multilocus introgression patterns: a case study on the mouse X chromosome in central Europe. Evolution, 65, 1428–1446. https://doi.org/10.1111/j.1558-5646.2011.01228.x

Pritchard, J. K., Stephens, M., & Donnelly, P. (2000). Inference of population structure using multilocus genotype data. Genetics, 155, 945–959.

Rousset, F., 2008. Genepop’007: a complete reimplementation of the Genepop software for Windows and Linux. Molecular Ecology Resources, 8, 103–106. https://doi.org/10.1111/j.1471-8286.2007.01931.x

Schoorl, J., & Zuiderwijk, A. (1981). Ecological isolation in *Triturus cristatus* and *Triturus marmoratus* (Amphibia: Salamandridae). Amphibia-Reptilia, 1, 235–252. https://doi.org/10.1163/156853881X00357

Scribner, K. T., & Avise, J. C. (1993). Cytonuclear genetic architecture in mosquitofish populations and the possible roles of introgressive hybridization. Molecular Ecology, 2, 139–149. https://doi.org/10.1111/j.1365-294X.1993.tb00103.x

Szymura, J. M., & Barton, N. H. (1986). Genetic analysis of a hybrid zone between the fire-bellied toads, *Bombina bombina* and *B. variegata*, near Cracow in southern Poland. Evolution, 40, 1141–1159. https://doi.org/10.1111/j.1558-5646.1986.tb05740.x

Trochet, A., Moulherat, S., Calvez, O., Stevens, V. M., Clobert, J., & Schmeller, D. S. (2014). A database of life-history traits of European amphibians. Biodiversity Data Journal, 2, e4123. https://doi.org/10.3897/BDJ2.e4123

Vallée, L. (1959). Recherches sur *Triturus blasii* de l’Isle: hybride naturel de *Triturus cristatus* Laur. X *Triturus marmoratus* Latr. Mémoires de la Société Zoologique de France, 31, 1–95.

Visser, M., de Leeuw, M., Zuiderwijk, A., & Arntzen, J. W. (2017). Stabilization of a salamander moving hybrid zone. Ecology and Evolution, 7, 689–696. https://doi.org/10.1002/ece3.2676

Wielstra, B. (2019). Historical hybrid zone movement: More pervasive than appreciated. Journal of Biogeography, 46, 1300–1305. https://doi.org/10.1111/jbi.13600

Wielstra, B., Burke, T., Butlin, R. K., & Arntzen, J. W. (2017a). A signature of dynamic biogeography: enclaves indicate past species replacement. Proceedings of the Royal Society B, Biological Sciences, 284, 20172014. https://doi.org/10.1098/rspb.2017.2014

Wielstra, B., Burke, T., Butlin, R. K., Avcı, A., Üzüm, N., Bozkurt, E., Olgun, K., & Arntzen, J. W. (2017b). A genomic footprint of hybrid zone movement in crested newts. Evolution Letters, 1, 93–101. https://doi.org/10.1002/evl3.9

Wielstra, B., Duijm, E., Lagler, P., Lammers, Y., Meilink, W. R., Ziermann, J. M., & Arntzen, J. W. (2014). Parallel tagged amplicon sequencing of transcriptome-based genetic markers for *Triturus* newts with the Ion Torrent next-generation sequencing platform. Molecular Ecology Resources, 14, 1080–1089. https://doi.org/10.1111/1755-0998.12242

Wielstra, B., McCartney-Melstad, E., Arntzen, J. W., Butlin, R. K., & Shaffer, H. B. (2019). Phylogenomics of the adaptive radiation of *Triturus* newts supports gradual ecological niche expansion towards an incrementally aquatic lifestyle. Molecular Phylogenetics and Evolution, 133, 120–127. https://doi.org/10.1016/j.ympev.2018.12.032

Zuiderwijk, A. (1990). Sexual strategies in the newts *Triturus cristatus* and *Triturus marmoratus*. Contributions to Zoology, 60, 51–64. https://doi.org/10.1163/26660644-06001003

